# RUNX1 suppresses breast cancer stemness and tumor growth

**DOI:** 10.1101/315093

**Authors:** Deli Hong, Andrew J. Fritz, Kristiaan H. Finstad, Mark P. Fitzgerald, Adam L. Viens, Jon Ramsey, Janet L. Stein, Jane B. Lian, Gary S. Stein

**Affiliations:** Department of Biochemistry and University of Vermont Cancer Center, University of Vermont College of Medicine, 89 Beaumont Avenue, Burlington, VT, 05405, USA.; Graduate Program in Cell Biology, University of Massachusetts Medical School, 55 Lake Avenue North, Worcester, MA 01655, USA

**Author notes:** **Corresponding author:** Gary S. Stein, Department of Biochemistry and University of Vermont Cancer Center, University of Vermont College of Medicine, 89 Beaumont Avenue, Burlington, VT 05405, USA, E.

## Abstract

Recent studies have revealed that mutations in the transcription factor Runx1 are prevalent in breast tumors. Yet, how loss of Runx1 contributes to breast cancer (BCa) remains unresolved. We demonstrate for the first time that Runx1 represses the breast cancer stem cell (BCSC) phenotype and consequently, functions as a tumor suppressor in breast cancer. Runx1 ectopic expression in MCF10AT1 and MCF10CA1a BCa cells reduces (60%) migration, invasion and *in vivo* tumor growth in mouse mammary fat pad (P<0.05). Runx1 is decreased in BCSCs, and overexpression of Runx1 suppresses tumorsphere formation and reduces the BCSC population. Furthermore, Runx1 inhibits Zeb1 expression, while Runx1 depletion activates Zeb1 and the epithelial-mesenchymal transition. Mechanistically Runx1 functions as a tumor suppressor in breast cancer through repression of cancer stem cell activity. This key regulation of BCSCs by Runx1 may be shared in other epithelial carcinomas, highlighting the importance of Runx1 in solid tumors.

## Introduction

Breast tumors are heterogeneous, as they are comprised of several types of cells, including transformed cancer cells, supportive cells, tumor-infiltrating cells and cancer stem cells (CSC). The CSC is acknowledged to be a significant component of growing tumors (1) (2). As the name implies, CSC can self-renew and reconstitute the cellular hierarchy within tumors (3) (4). Moreover, these stem-like cells are highly chemo-resistant and metastatic (5) (6). Significantly, signaling pathways (TGF-β, WNT, Hedgehog and Notch) and transcription factors (Snail, Twist and Zeb) regulating stemness properties in CSC are involved in controlling an essential cellular process designated epithelial-mesenchymal transition (EMT) (7, 8). The EMT process is linked to chemo-resistance and cancer metastasis (9, 10) (11). One such example is Zeb1, a well-known EMT-activator that is also a key factor for cell plasticity and promotes stemness properties in breast and pancreatic cancers (12) (13). However there remains a compelling requirement to understand regulatory mechanisms that contribute to and sustain the stemness of the CSC population. By identifying regulator(s) that maintain or repress the cancer stem cell phenotype can provide insights for novel therapeutic approaches. Recently, a list of 40 mutation-driver genes for which deregulation contributes directly to breast tumor progression has been identified (14); among these is the transcription factor RUNX1, which has been shown to repress EMT. Here we address for the first time, the function of RUNX1 in regulating breast cancer stem cells.

The Runx family, including RUNX1, Runx2 and Runx3, are evolutionarily conserved transcription factors and function as critical lineage determinants of various tissues (15).

During normal development, it is well documented that RUNX1 plays a fundamental role in controlling the stem cell population in hematopoietic (16) (17) (18), hair follicle (19) (20), gastric (21) and oral epithelial stem cells (22). As a master transcriptional regulator, RUNX1 is a central player in fine-tuning the balance among cell differentiation, proliferation, and cell cycle control in stem cells during normal development (23). In the mammary gland, it has recently been shown that RUNX1 is involved in luminal development (24). These studies also showed that loss of RUNX1 in mammary epithelial cells blocked differentiation into ductal and lobular tissues. These findings suggest that RUNX1 is an essential regulator of normal mammary stem cells (24). In addition to its essential function during normal development, disrupting RUNX1 function(s) can cause cancer (25) (15). RUNX1 is a frequent target of translocations and other mutations in hematopoietic malignancies. For example, RUNX1 related chromosomal translocations including RUNX1-ETO (26), TEL-RUNX1 (27) and RUNX1-EVI (28) are associated with distinct leukemia subtypes.

In breast cancer, RUNX1 has been shown to regulate the WNT pathway and key transcription factors including ERα and Elf5 (15) (29) (30) (31). Recent studies from our group have demonstrated that RUNX1 has tumor suppressor activity by maintaining the epithelial phenotype and repressing EMT (32). RUNX1 expression is decreased during breast cell EMT, and loss of RUNX1 expression in normal-like epithelial cells (MCF10A) and epithelial like breast cancer cells (MCF7) initiates the EMT process. Complementary studies demonstrated that ectopic expression of RUNX1 reverses cells to the epithelial state. However mechanisms underlying RUNX1 regulation of cancer stem cell properties and the consequences for tumor growth *in vivo* remain to be resolved.

Based on evidence that RUNX1 regulates stem cell properties during normal development and that loss of RUNX1 activates partial EMT in breast cancer, we hypothesized that RUNX1 represses the cancer stem cell population and/or stemness properties in breast cancer. We investigated whether altering RUNX1 levels by overexpression and knockdown in breast cancer cells changes the stemness phenotype, aggressive properties and tumor progression *in vivo*. Our findings have identified for the first time a significant function for RUNX1 in repressing the cancer stem cell population as well as tumorsphere formation, and demonstrated that RUNX1 represses breast cancer tumor growth *in vivo*.

## Materials and Methods

### Cell culture

MCF10AT1 and MCF10A cells were grown in DMEM: F12 (Hyclone: SH30271, Thermo Fisher Scientific, Waltham, MA) with 5% (v/v) horse serum (Gibco: 16050, Thermo Fisher Scientific, Waltham, MA, USA) + 10 μg/ml human insulin (Sigma Aldrich, St. Louis, MO: I-1882) + 20 ng/ml recombinant hEGF (Peprotech, Rocky Hill, NJ, USA: AF-100-15) + 100 ng/ml cholera toxin (Sigma Aldrich: C-8052) + 0.5 μg/ml hydrocortisone (Sigma Aldrich: H-0888) 50 IU/ml penicillin/50 μg/ml streptomycin and 2 mM glutamine (Life Technologies, Carlsbad, CA, USA: 15140-122 and 25030-081, respectively). MCF10CA1a cells were grown in DMEM: F with 12, 5% (v/v) horse serum with 50 IU/ml penicillin/50 μg/ml streptomycin and 2 mM glutamine. MCF7 cells were maintained in Dulbecco modified Eagle medium (DMEM) high glucose (Fisher Scientific: Thermo Fisher Scientific, Waltham, MA, USA: MT-10-017-CM) supplemented with 10% (v/v) FBS (Atlanta Biologicals, Flowery Branch, GA, USA: S11550), 50 IU/ml penicillin/50 μg/ml streptomycin.

### Lentiviral plasmid preparation and viral vector production

RUNX1 cDNA was cloned into Lentivirus-based overexpression plasmids pLenti-CMV-Blast-DEST (Addgene). To generate lentivirus vectors, 293T cells in 10 cm culture dishes were co-transfected with 10 μg of pGIPZ shRUNX1 or pGIPZ non-silencing, with 5 μg of psPAX2, and 5 μg of pMD2.G using lipofectamine 2000 reagent (Life Technologies). Viruses were harvested every 48 h post-transfection. After filtration through a 0.45 μm-pore-size filter, viruses were concentrated by using LentiX concentrator (Clontech, Mountain View, CA, USA).

### Gene delivery by transfection and infection

For overexpression RUNX1, MCF10AT1 or MCF10CA1a cells were plated in six-well plates (1x10^5^ cells per well) and infected 24 h later with lentivirus expressing RUNX1 overexpression or Empty Vector. Briefly, cells were treated with 0.5 ml of lentivirus and 1.5 ml complete fresh DMEM-F12 per well with a final concentration of 4 μg/ml polybrene. Plates were centrifuged upon addition of the virus at 1460 × *g* at 37°C for 30 min. Infection efficiency was monitored by GFP co-expression at 2 days post infection. Cells were selected with 2 μg/ml puromycin (Sigma Aldrich P7255-100MG) for at least two additional days. After removal of the floating cells, the remaining attached cells were passed and analyzed. ShRUNX1 virus were generated and delivered as has been described previously (32).

### Western blotting

Cells were lysed in RIPA buffer and 2X SDS sample buffer supplemented with cOmplete, EDTA-free protease inhibitors (Roche Diagnostics) and MG132 (EMD Millipore San Diego, CA, USA). Lysates were fractionated in an 8.5% acrylamide gel and subjected to immunoblotting. The gels are transferred to PVDF membranes (EMD Millipore) using a wet transfer apparatus (Bio-Rad Laboratories, Hercules, CA, USA). Membranes were blocked using 5% Blotting Grade Blocker Non-Fat Dry Milk (Bio-Rad Laboratories) and incubated overnight at 4°C with the following primary antibodies: a rabbit polyclonal RUNX1 (Cell Signaling Technology, Danvers, MA, USA: #4334, 1:1000); a mouse monoclonal to E-cadherin (Santa Cruz Biotechnology, Inc., Santa Cruz, CA, USA: sc21791, 1:1000); a mouse monoclonal Vimentin (Santa-Cruz Biotechnology sc-6260, 1:1000); a mouse monoclonal to β-Actin (Cell Signaling Technology #3700, 1:1000); a rabbit polyclonal Twist1 (Santa Cruz Biotechnology sc-15393, 1:2000); a rabbit polyclonal Zeb1 (Sigma-Aldrich HPA027524-100UL, 1:1000). Secondary antibodies conjugated to HRP (Santa Cruz Biotechnology) were used for immunodetection, along with the Clarity Western ECL Substrate (Bio-Rad Laboratories) on a Chemidoc XRS+ imaging system (Bio-Rad Laboratories).

### Tumorsphere formation assay

Monolayer cells were enzymatically dissociated into single cells with 0.05% trypsin-EDTA. Cells were plated at 10,000 cells per well in a 24-well low-attachment plate (Corning). Cells were grown for 7 days in DMEM/F12 supplemented with B27 (Invitrogen) in the presence of 10 ng/ml EGF and 10 ng/ml bFGF. Where indicated, the CDK4 inhibitor palbocilib (Sigma) was added at a final concentration of 100 nM. Tumorsphere-forming efficiency was calculated as the number of spheres divided by the number of singles cells seeded, expressed as a percentage.

### CD24/CD44 flow cytometry

Flow cytometry for CD24 (PE-cy7, Biolegend 311120) and CD44 (APC, BD Pharmigen 559942) was performed using the best conditions for marker detection as previously described (33) (34). Cells were grown to sub-confluency and dissociated with Accutase. The Accutase was quickly neutralized with serum and 1x10^6^ cells were washed with 1xPBS. These cells were then re-suspended in 475ul of 1%FBS/ 1xPBS, to which 25ul of CD44-APC and 4ul of CD24-PE-cy7 were added and incubated at room temperature for 30 minutes. Cells were then washed with PBS and strained (Falcon 352235) to obtain single cell suspensions. Isotype controls were used to gate for negative signal in each replicate of the experiment.

### Migration assays

For the scratch assays, cells were seeded in triplicate and when they reached 95–100% confluence, were serum starved with 0.1% FBS-containing media for 12 h. Subsequently, a scratch was made across the cell layer using a P-200 pipette tip, and cell migration was monitored by recording images at indicated time points post-scratch. The area of the scratch was quantified using the MiToBo plug-in for ImageJ software and plotted as a percentage of total area.

For the transwell migration assay, cells were trypsinized and re-seeded in triplicate in migration chambers (BD Bioscience, Bedford, MA) in serum-free medium. 24 hours (MCF10AT1 cells) or 48 hours (MCF10CA1a cells) after cell seeding, the experiment was performed and results quantified as previously described (35).

### Invasion Assay

For the invasion assay, cells were trypsinized and reseeded in triplicate in growth factor-reduced Matrigel invasion chambers (BD Bioscience, Bedford, MA) in serum-free medium. 24 hours (MCF10AT1 cells) or 48 hours (MCF10CA1a cells) after cell seeding, the experiment was performed and results quantified as previously described (35).

### Immunofluorescence staining microscopy

Cells grown on coverslips were fixed with using 3.7% formaldehyde in Phosphate Buffered Saline (PBS) for 10 min. Cells were then permeabilized in 0.1% Triton X-100 in PBS, and washed in 0.5% Bovine Serum Albumin in PBS. Detection was performed using a rabbit polyclonal RUNX1 antibody (Cell Signaling #4336), a mouse monoclonal CD24 (Santa-Cruz sc-11406). Staining was performed using fluorescent secondary antibodies; for rabbit polyclonal antibodies a goat anti-rabbit IgG (H+L) secondary antibody, Alexa Fluor^®^ 568 conjugate (Life Technologies A-11011), was used and for mouse monoclonal a F(ab')2-goat anti-mouse IgG (H+L) secondary antibody, Alexa Fluor^®^ 488 conjugate was used (Life Technologies A-11001).

### Animal studies

Female SCID mice 7 weeks of age were used for mammary fat pad injection. The mice were randomly divided into two groups (seven for each group). In all, 1X10^6^ MCF10CA1a cells suspended in 0.1 ml of saline were mixed with 0.1 ml of Matrigel (BD) and were injected under mammary fat pads. Bioluminescence images were acquired by using the IVIS Imaging System (Xenogen) 5 min after injection 150 mg/kg of D-Luciferin (Gold BioTech, St. Louis, MO) in PBS. All animals were housed in a pathogen-free environment and handled according to protocol number 12-051 approved by the Institutional Animal Care and Use Committee at the University of Vermont. In conducting using animals, the investigators adhere to the laws of the United States and regulations of the Department of Agriculture.

#### Analysis of RUNX1 expression and patient survival using public data sets

The PROGgene database (www.compbio.iupui.edu/proggene) (36) (37) was used to identify the data sets for survival analysis and re-analyzed the public GEO data sets (www.ncbi.nlm.nih.gov/gds) (GSE37751 (38), GSE7390 (39), TCGA (40)). RUNX1 expression in different breast cancer stages was analyzed using the TCGA database (www.cbioportal.org).

### Quantitative PCR

RNA was isolated with Trizol (Life Technologies) and cleaned by DNase digestion (Zymo Research, Irvine, CA, USA). RNA was reversed transcribed using SuperScript II and random hexamers (Life Technologies). cDNA was then subjected to quantitative PCR using SYBR Green technology (Applied Biosystems, Foster City, CA, USA).

RUNX1 Forward: AACCCTCAGCCTCAGAGTCA,
RUNX1 Reverse: CAATGGATCCCAGGTATTGG;
FN1 Forward: CATGAAGGGGGTCAGTCCTA;
FN1 Reverse: CTTCTCAGCTATGGGCTTGC;
VEGF Forward: CCTTGCTGCTCTACCTCCAC;
VEGF Reverse: CCATGAACTTCACCACTTCG;
CXCR4 Forward: TACACCGAGGAAATGGGCTCA;
CXCR4 Reverse: TTCTTCACGGAAACAGGGTTC;
CXCL12 Forward: GTGGTCGTGCTGGTCCTC;
CXCL12 Reverse: AGATGCTTGACGTTGGCTCT;
MMP13 Forward: ATGAGCCAGAGTGTCGGTTC;
MMP13 Reverse: GTTAGTAGCGACGAGCAGGAC;
MMP9 Forward: ATAGACTACTACAGGCT;
MMP9 Reverse: TAGCACGGATAGACCA;
GAPDH Forward: TGTGGTCATGAGTCCTTCCA,
GAPDH Reverse: ATGTTCGTCATGGGTGTGAA;
HPRT Forward: TGCTGACCTGCTGGATTACA,
HPRT Reverse: TCCCCTGTTGACTGGTCATT;
β-Actin Forward: AGCACAGAGCCTCGCCTTT,
β-Actin Reverse: CGGCGATATCATCATCCAT.

### ChIP-qPCR

ChIP-qPCR was performed essentially as described (41). Briefly, 200,000 MCF10AT1 or MCF10CA1a cells were cross-linked, lysed and sonicated to obtain DNA fragments mostly in the 200-1000-bp range. Immunoprecipitation was performed at 4°C overnight with anti-RUNX1 antibody (4334, Cell Signaling Technology) at a 1:15 antibody to chromatin ratio. Primers used in ChIP-qPCR are listed below:

Zeb1 Forward: GTCGTAAAGCCGGGAGTGTC,
Zeb1 Reverse: GCCATCCGCCATGATCCTC;
ZNF333 (negative control 1) Forward: TGAAGACACATCTGCGAACC,
ZNF333 Reverse: TCGCGCACTCATACAGTTTC;
ZNF180 (negative control 2) Forward: TGATGCACAATAAGTCGAGCA,
ZNF180 Reverse: TGCAGTCAATGTGGGAAGTC.

### Tissue microarray

Tissue microarray data of RUNX1 in breast cancer patients were obtained from Human Protein Atlas (www.proteinatlas.org) (42).

### Statistical analysis

Each experiment was repeated at least three times. The differences in mean values among groups were evaluated and expressed as the mean ± SEM. A *P*-value less than 0.05 was considered statistically significant (^*^*P* < 0.05, ^**^*P* < 0.01, ^***^*P* < 0.001). Student’s *t*-test was used to compare the expressions of cell surface markers, side population analysis, cell viability, relative mRNA levels, migrated cells and invaded cells.

## Results

### 1. Reduced RUNX1 expression is associated with decreased survival probability in breast cancer patients

To investigate possible association between RUNX1 expression and breast cancer progression, we first examined RUNX1 expression in normal and breast cancer patients using the Human Protein Atlas. Within normal breast tissues, RUNX1 is highly expressed in the mammary gland (Fig. 1A). However in ductal carcinoma tissues, the level of RUNX1 is decreased in malignant regions (red circle) compared with normal glandular tissues (blue circle) in the same tumor specimen (Fig. 1B). In the majority of ductal carcinoma specimens (9 out 12 samples) from the Human Protein Atlas, 75% of breast cancer tumors show low RUNX1 staining (Fig. 1C). We also analyzed TCGA data and found that RUNX1 levels are progressively decreased across early stage breast cancer (Stage 1 vs Stage2; Stage 2 vs Stage 3) (Supplemental Figure 1). These findings suggest that during breast cancer progression, the mammary gland loses its original structure and RUNX1 levels are decreased. The data are consistent with our previous report that RUNX1 is highly expressed in normal-like mammary epithelial MCF10A cells and reduced in a panel of breast cancer cell lines (32). With the reduced RUNX1 expression, mammary epithelial cells do not maintain their epithelial phenotype (32) From these observations of low RUNX1 in breast tumors and the concomitants loss of RUNX1 in normal epithelial cells with loss of epithelial properties, we hypothesized that loss of RUNX1 is promoting a breast cancer phenotype.

**Figure 1.**
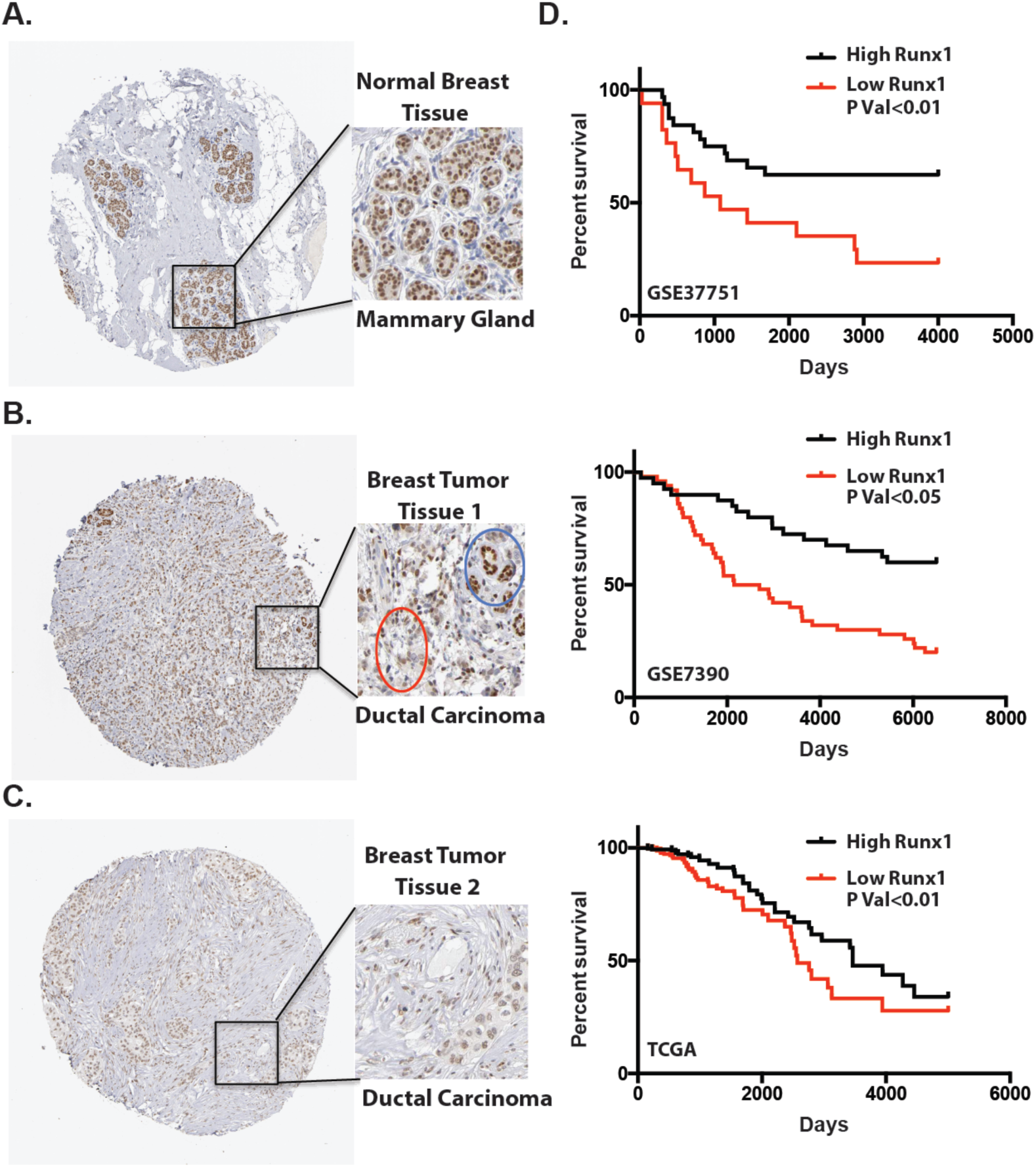
Reduced RUNX1 expression is associated with decreased survival probability in breast cancer patients. **(A)** Representative tissue microarray images of RUNX1 in normal breast tissue. **(B)** and **(C)** Representative tissue microarray images of RUNX1 in breast tumor tissues. **(D)** Kaplan-Meier analysis showed higher overall survival in patients with higher RUNX1 mRNA expression (GSE37751, GSE7390 and TCGA). Gehan-Breslow-Wilcoxon test with *p* value<0.01, *p* value<0.05, *p* value<0.01 respectively compared with high RUNX1 expression patients and low RUNX1 expression patients in three data sets.

We therefore addressed whether there was a clinical relation of RUNX1 expression in breast cancer patient tumors to survival. Using publically available mRNA expression datasets, we analyzed the correlation of mean expression levels of RUNX1 and survival rate in breast cancer patient tissue samples. Kaplan–Meier analysis of the expression of RUNX1 in three separate datasets of GSE37751- “Molecular Profiles of Human Breast Cancer and Their Association with Tumor Subtypes and Disease Prognosis” (36 high RUNX1 and 24 low RUNX1 patients), GSE7390-“Strong Time Dependence of the 76-Gene Prognostic Signature” (82 high RUNX1 and 116 low RUNX1 patients) and TCGA data of breast cancer patients mRNAs (304 high RUNX1 and 290 low RUNX1 patients) indicated a statistically significant correlation (p < 0.01, p < 0.05, and p<0.01 respectively) between high RUNX1 expression levels and longer patient survival time (Fig. 1D). These results suggested that reduction in RUNX1 expression is associated with low survival probability of breast cancer patients. Thus several *in vitro* studies combined with these clinical observations support a role for RUNX1 in repressing tumor growth.

### 2. RUNX1 is decreased in tumors formed in mouse mammary fat pad

To further establish if RUNX1 decreases during breast tumor growth *in vivo*, we utilized a mouse xenograft model to examine RUNX1 levels before and after tumor formation. MCF10CA1a cells, which are aggressive breast cancer cells, were injected into mammary fat pad of SCID mice and tumor growth was monitored weekly. Tumors formed within two weeks (Fig. 2A), and one month post-injection, mice were sacrificed and tumors were removed to analyze for RUNX1 and other factors at both protein and mRNA levels. The parental MCF10CA1a cells had a 3.3 fold higher RUNX1 protein level than the removed tumor (Fig. 2B,C). Q-PCR using human-specific primer sets confirmed that RUNX1 mRNA is also decreased specifically within the tumor (Fig. 2C). The epithelial marker E-cadherin was decreased in tumor samples, while the mesenchymal marker Vimentin was increased (Fig. 2B). In addition to Vimentin, the mRNA levels of several human cancer-related genes such as VEGF, FN1, MMP13, MMP9, CXCR4, CXCL12 are also up regulated (Fig. 2B, 2D). These findings indicate that the human breast cancer cells that formed a tumor in mouse mammary fat pads acquired a more aggressive phenotype and that RUNX1 expression is decreased during the period of tumor growth. Therefore we have directly demonstrated that in this MCF10CA1a mouse xenograft model, RUNX1 expression is decreased during *in vivo* model of tumor progression.

**Figure 2.**
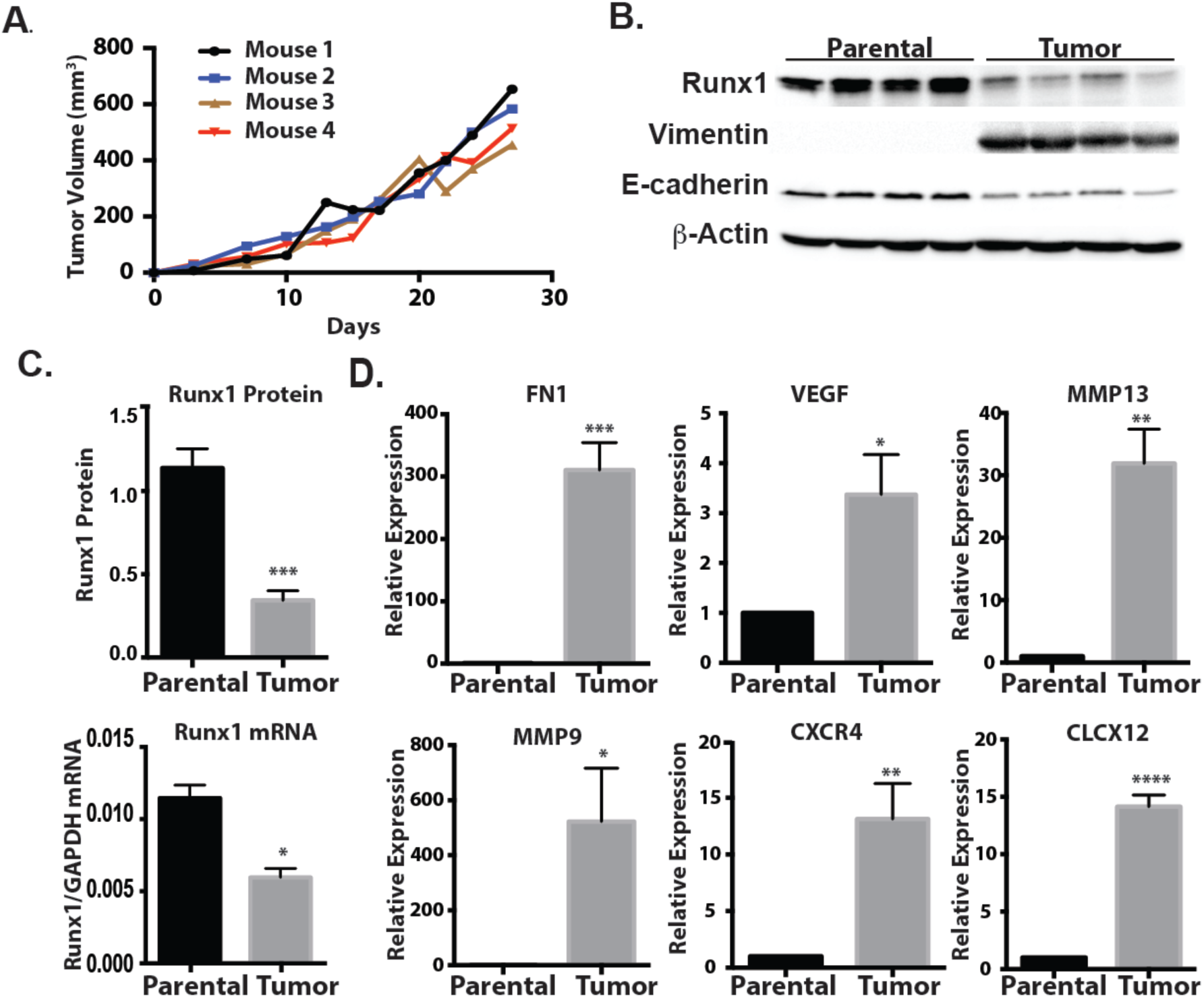
RUNX1 is decreased in tumors formed in mouse mammary fat pad. **(A)** MCF10CA1a cells were injected into the mammary fat pad of SCID mice. Points represent mean tumor volume. **(B)** Western blot analyses show RUNX1 and E-cadherin levels are decreased and Vimentin level is increased in tumor samples compared to MCF10CA1a cells. **(C)** Upper panel, Protein quantification show that RUNX1 is significant decreased in tumor samples compared to MCF10CA1a. Data shown represent mean ± SEM from three independent experiments. Lower panel, RT-qPCR analyses of RNA from tumor samples show decreased RUNX1 expression of compared with MCF10CA1a cells. Student’s *t* test ^*^ *p* value <0.05, ^***^ *p* value <0.001 and. Error bars represent the standard error of the mean (SEM) from three independent experiments. **(D)** RT-qPCR analyses of RNA from tumor samples show activation of mesenchymal marks Vimentin and FN1 and other tumor growth related genes including MMP9, MMP13, VGF, CXCR4 and CXCL12 compared with MCF10CA1a cells. Student’s *t* test ^*^ *p* value <0.05, ^**^ p value <0.01, ^***^ *p* value <0.001 and ^****^ *p* value <0.0001. Error bars represent the standard error of the mean (SEM) from three independent experiments.

### 3. RUNX1 reduces the aggressive phenotype of breast cancer cells *in vitro*

It has been suggested that RUNX1 has tumor suppressor activity (29, 30, 32). Based on these data and the results that RUNX1 level is decreased in the xenograft model (Fig. 2B), we further addressed whether ectopic expression of RUNX1 in malignant breast cancer cells reduces the aggressive phenotype. RUNX1 was overexpressed using a lentivirus delivery system (pLenti-CMV) in pre-malignant MCF10AT1 and highly aggressive malignant MCF10Ca1a cells (Fig. 3A). Upon overexpressing RUNX1, Vimentin expression is decreased in both cell lines (Fig. 3A). However, E-cadherin expression was not affected by RUNX1 overexpression, suggesting that the cells have not fully transitioned back to normal-like stage.

**Figure 3.**
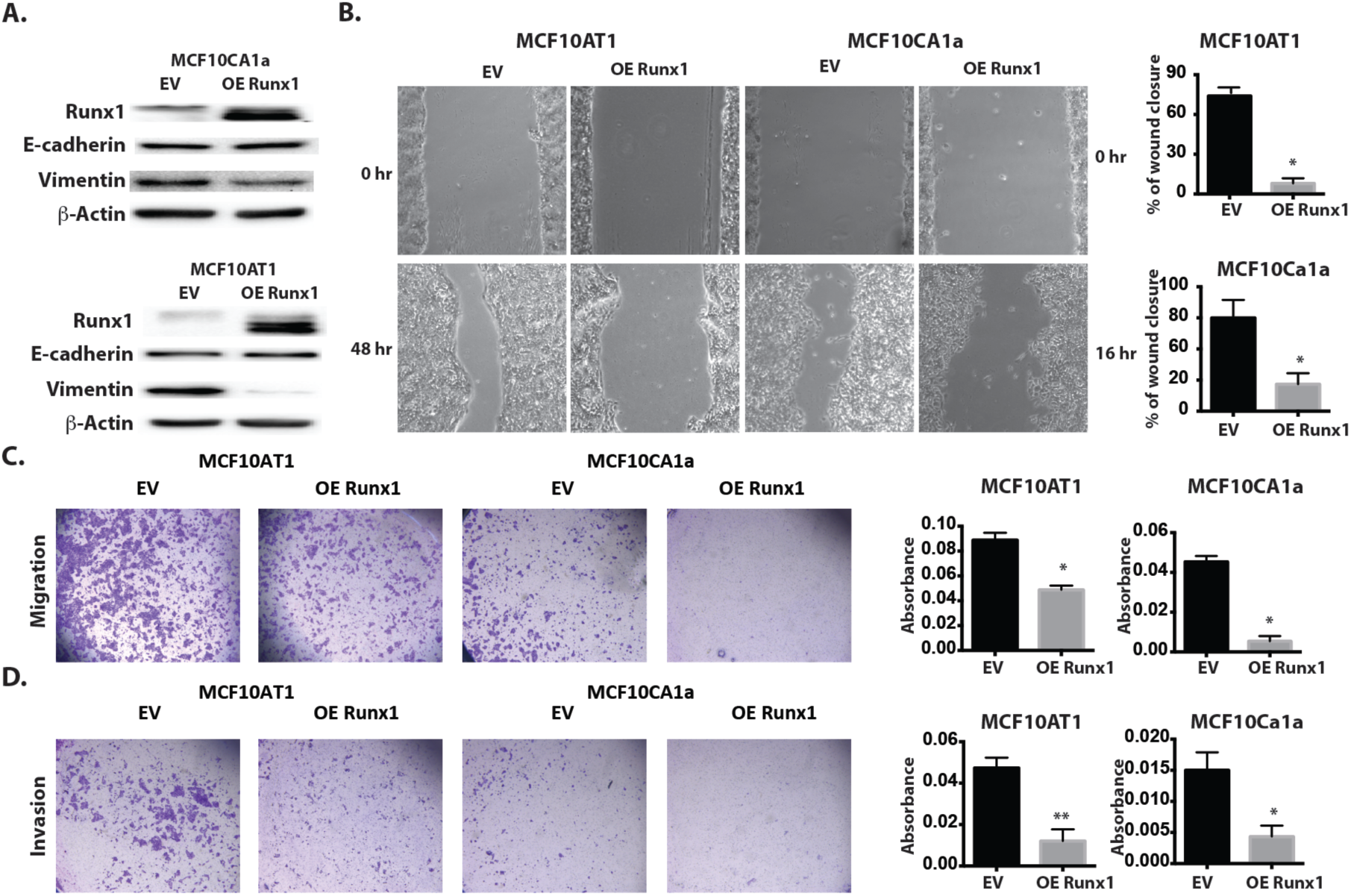
RUNX1 reduces the aggressive phenotype of breast cancer cells *in vitro.* **(A)** Western blot analyses confirm RUNX1 overexpression in MCF10CA1a (Upper) and MCF10AT1 (Lower) cells. Vimentin expression is repressed upon RUNX1 overexpression in both cell lines. **(B)** Representative phase contrast images (magnification 100×) of MCF10AT1 and MCF10CA1a cells with EV control or RUNX1 overexpression subjected to a scratch assay for times indicated. The area of the scratch was plotted as a percentage of total area for N = 3 independent experiments carried out in duplicate. Data shown represent mean ± SEM from three independent experiments using student t-test. **(C)** Light microscopy images (mag. 12×) of stained cells from a representative (1 of N = 2) trans-well migration assay experiment MCF10AT1 and MCF10CA1a cells with EV control or RUNX1 overexpression (*left*); quantitation of migrated cells assessed by measurement of the absorbance of solubilized crystal violet stain retained by migrated cells (*right*). **(D)** Light microscopy images (mag. 12×) of stained cells from a representative (1 of N = 2) trans-well matrigel invasion assay experiment with MCF10AT1 and MCF10CA1a cells with EV control or RUNX1 overexpression to evaluate invasion (*left*); quantitation of invaded cells assessed by measurement of the absorbance of solubilized crystal violet stain retained by invaded cells (*right*). For all assays, three independent experiments were carried out in duplicates. All quantitative data are depicted as mean ± S.E.M per group. ^*^*P* < 0.05, ^**^*P* < 0.01 (student’s *t*-test).

To evaluate the effect of RUNX1 in regulation of migration and invasion capacities of the breast cancer cells *in vitro,* we used the scratch migration and Transwell assays. Figure 3B shows representative images of the scratch assay, both at the time of the scratch and 48 h (MCF10AT1) or 16 h (MCF10CA1a) later. RUNX1 overexpression decreases the ability of breast cancer cells to migrate. These results were confirmed using the trans-well migration assay (Fig. 3C). Invasion of both MCF10AT1 and MCF10CA1a cells was also significantly inhibited when RUNX1 was overexpressed (Fig. 3D). We conclude from these studies that loss of RUNX1 in MCF10A and cancer cells is detrimental in promoting EMT in vitro(32) and *in vivo* (Fig 2B), while exogenous expression of RUNX1 suppresses the migration and invasion of breast cancer cells *in vitro*.

### 4. RUNX1 represses tumor growth *in vivo*

Together our data above and the earlier studies demonstrate that RUNX1 has tumor suppresser activity *in vitro.* However, to date there are no studies showing that RUNX1 inhibits tumor growth *in vivo*. We tested the ability of RUNX1 to alter tumor growth *in vivo* by using the metastatic MCF10CA1a breast cancer cells. MCF10CA1a/EV (control) and MCF10CA1a/ RUNX1-overexpression cells carrying a luciferase reporter (experiment) were injected into the mammary fat pad of SCID mice. Eighteen days post-injection tumors appeared in the control mice, with an average volume of 63 mm^3^ (caliper measurement), while the experimental group had barely palpable tumors (Fig. 4A). At the end point of this experiment (4 weeks), we sacrificed the mice, excised the tumors, and measured tumor volume and weight (Fig. 4B, C). Mice injected with MCF10CA1a/OE RUNX1 cells had a significantly reduced tumor size (57%) and weight (47%) compared with tumors from control mice. Supplemental Figures 2A and 2B show the excised tumors and luminescence of tumors in all seven mice from each group. MCF10CA1a cells with EV or OE RUNX1 were validated before injection into the SCID mice (Supplement Figure 2C). Luminescent images of representative mice (Fig. 4D) confirm reduced tumor growth. Collectively, these data indicate that RUNX1 inhibits tumorigenesis and suppresses breast cancer *in vivo*.

**Figure 4.**
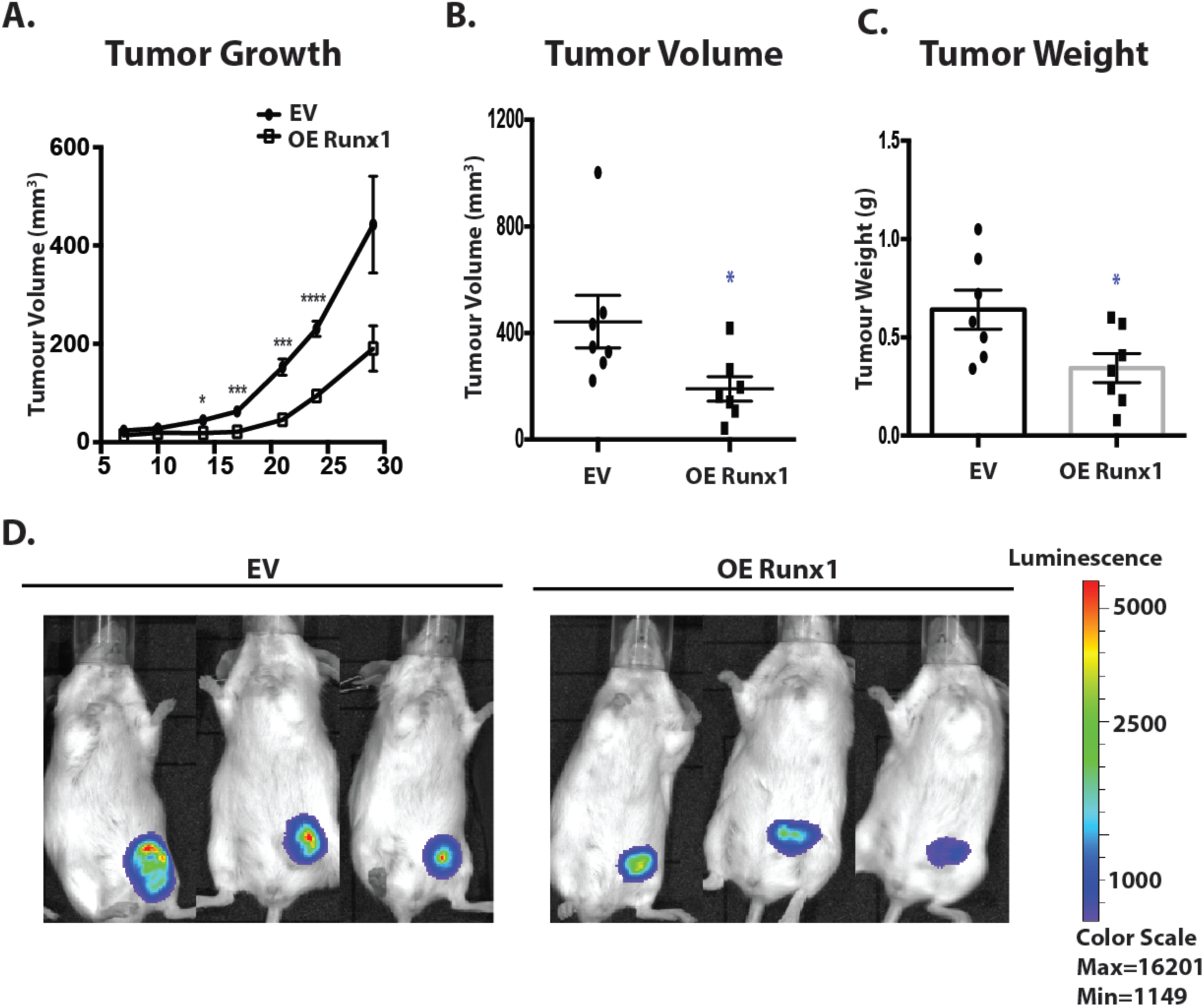
RUNX1 represses tumor growth *in vivo.* **(A)** A total of 1 × 10^6^ MCF10CA cells with EV or RUNX1 overexpression were injected into mammary fat pad of SCID mice (*n* = 7 in each group). The points represent average tumor volume at each time point ± S.E.M. *P* values were obtained by 2-tailed Student *t* test. ^*^, *P* < 0.05; ^***^, P<0.001; ^****^, P<0.0001. **(B)** Tumor size measured at day 28 (end point). *P* values were obtained by 2-tailed Student *t* test. ^*^, *P* < 0.05. **(C)** Tumor weight at day 28 (end point). *P* values were obtained by 2-tailed Student *t* test. ^*^, *P* < 0.05. **(D)** Representative luminescence images at 4 weeks after mammary fat pad injection.

### 5. RUNX1 level is decreased in breast cancer stem cells (BCSC)

As breast cancer stem cells have been shown to be critical for tumor initiation and growth (11) and all of our data demonstrate a role for RUNX1 in decreasing tumorigenesis, we next investigated the potential role of RUNX1 in breast cancer stemness. We used fluorescence-activated cell sorting (FACS) to isolate BCSCs from pre-malignant MCF10AT1 cells based on expression of the cell-surface antigen markers CD44 and CD24. These two markers have been successfully used to identify putative CSCs in primary breast tumors or mammary cell lines (CD44^high^/CD24^low^). We compared the BCSC cells with bulk cells (CD44^high^/CD24^high^) as gated in Supplement Figure 3A. The CD44^high^/CD24^low^ subpopulation from MCF10AT1 cells displayed lower levels of RUNX1 protein (33%) compared to the bulk cell population and the parental MCF10AT1 cells (Fig. 5A). To examine whether CD24^low^ cells have low RUNX1 expression, we also performed immunofluorescence co-staining of RUNX1 and CD24 in MCF10AT1 cells. The cells with high CD24 expression also have high RUNX1 expression (Supplement Figure 4). Moreover, the CD44^high^/CD24^low^ population displays many CSC-like properties; they are endowed with higher expression of cancer stem cell markers Zeb1 and Twist1 (Fig. 5A) and greater long-term self-renewal capacity as measured by tumorsphere formation assays (Fig. 5B). Collectively, these data provide evidence that cell populations with BCSC properties express lower levels of RUNX1 compared to the bulk and parental population, and suggest that RUNX1 influences BCSC properties.

**Figure 5.**
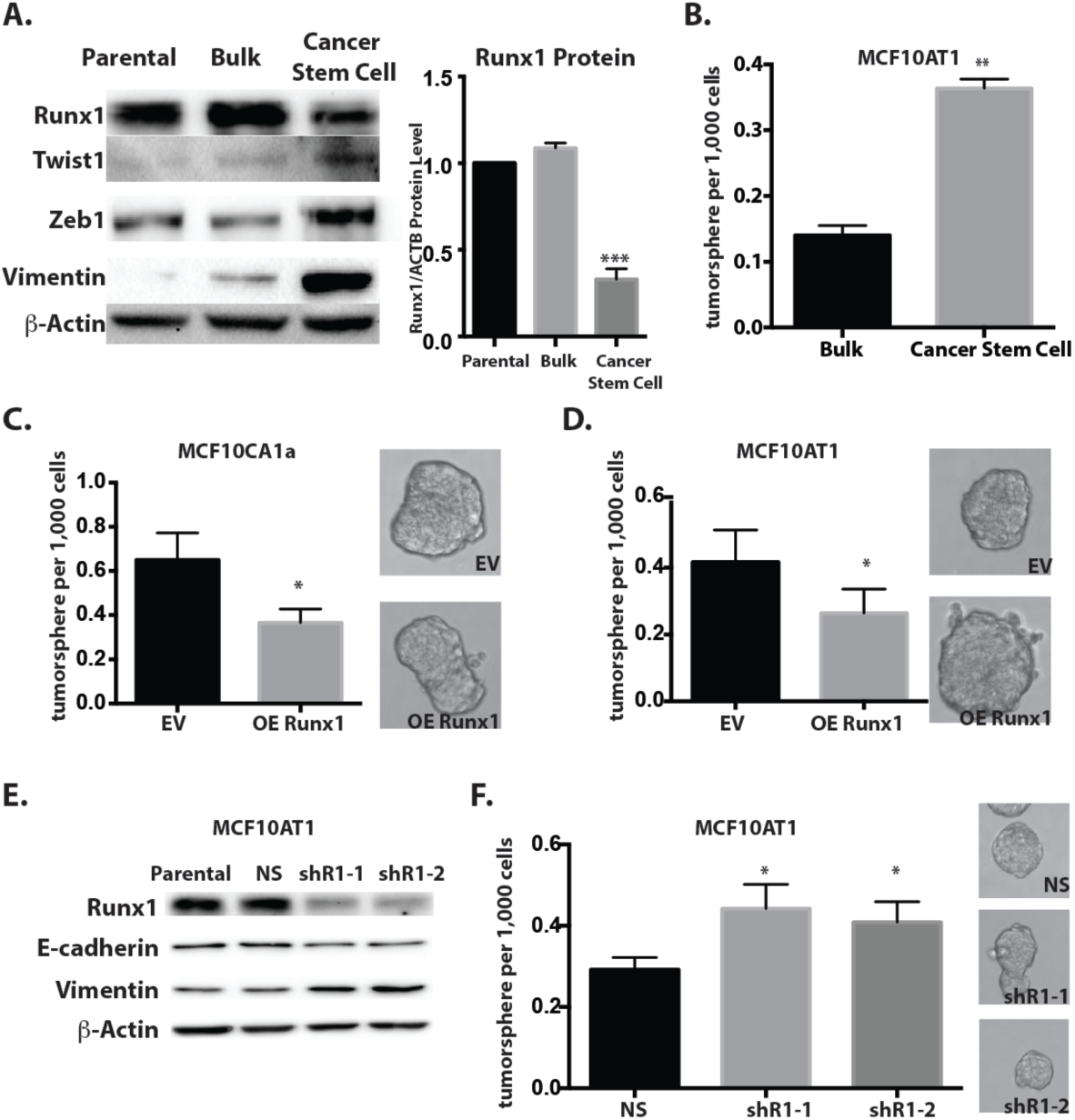
RUNX1 level is decreased in BCSC. **(A)** Western blot analyses show RUNX1 is decreased and Zeb1, Twist1 and Vimentin level are increased in BCSC samples compared to Parental and Bulk MCF10AT1 cells. Right, protein quantification shows that RUNX1 is significant decreased in BCSC. **(B)** Tumorsphere formation efficiency for BCSC populations is significantly higher than bulk population. Data shown represent mean ± SEM from three independent experiments. ^**^*P* < 0.01. **(C)** RUNX1 overexpression in MCF10CA1a cells reduces tumorsphere formation efficiency. Data shown represent mean ± SEM from three independent experiments. ^*^*P* < 0.05. Right, represent picture of tumorsphere. **(D)** RUNX1 overexpression in MCF10AT1 cells reduces tumorsphere formation efficiency. Data shown represent mean ± SEM from three independent experiments. ^*^*P* < 0.05 Right, represent picture of tumorsphere. **(E)** Western blot analyses of lysates from MCF10AT1 cells treated with shRUNX1 show decreased protein expression of RUNX1 and E-cadherin and increased protein expression of Vimentin. **(F)** RUNX1 knockdown in MCF10AT1 cells activates tumorsphere formation efficiency. Data shown represent mean ± SEM from three independent experiments. ^*^*P* < 0.05. Right, represents picture of tumorsphere.

### 6. RUNX1 inhibits stemness properties in breast cancer cells

To further investigate the role of RUNX1 in regulating BCSC properties, we addressed the capability of RUNX1 to regulate tumorsphere formation from breast cancer cells. Tumorsphere formation assays were performed using non-adherent plates with non-serum medium. The ectopic expression of RUNX1 in both MCF10CA1a and MCF10AT1 cells significantly decreased the number of tumorspheres (p < 0.05) (Fig. 5C, 5D). To better understand if RUNX1 represses stemness properties in breast cancer, we used two lenti-viruses to establish RUNX1 knockdown cell lines in MCF10AT1 cells (Fig. 5E). Depletion of RUNX1 in these cell lines activated an epithelial to mesenchymal transition with lower E-cadherin and higher Vimentin expression (Fig. 5E). Significantly, the knockdown of RUNX1 resulted in increased tumorsphere formation efficiency in MCF10AT1 cells (51% and 41% respectively) (Fig. 5F). This ability of RUNX1 to repress stemness properties was also observed in additional cell lines, including normal-like MCF10A cells and ER positive luminal-like MCF7 cells (Supplement Figure 5A, B), which suggests that RUNX1 suppression of stemness is a universal phenotype in breast cancer cells.

Further evidence for the influence of RUNX1 on the cancer stem cell population in MCF10AT1 cells was provided by flow cytometry analysis. As shown in Figure 6A, ectopic expression of RUNX1 reduced the CD44^high^/CD24^low^ subpopulation of MCF10AT1 cells from 22.3% to 15.1% (Fig. 6A). Consistent with the consequence of RUNX1 overexpression, knockdown of RUNX1 significantly increased the CD44^high^/CD24^low^ subpopulation of MCF10AT1 cells by more than two fold (21.9% ns; 45.3% shR1-1; 45.6% shR1-2) (Fig. 6B). However ectopic expression of RUNX1 in MCF10CA1a cells did not change the percent of the CD44^high^/CD24^low^ cancer stem cell population (Supplement Figure 3B). The highly metastatic MCF10CA1a cells have a large percentage of cells (80%) that are CD44^high^/CD24^low^, indicating that the cells may have lost their plasticity and are locked into a mesenchymal phenotype (Supplement Figure 3B). These results indicate that RUNX1 functions both to suppress cancer stem cell properties and reduce the breast cancer stem cell population.

**Figure 6.**
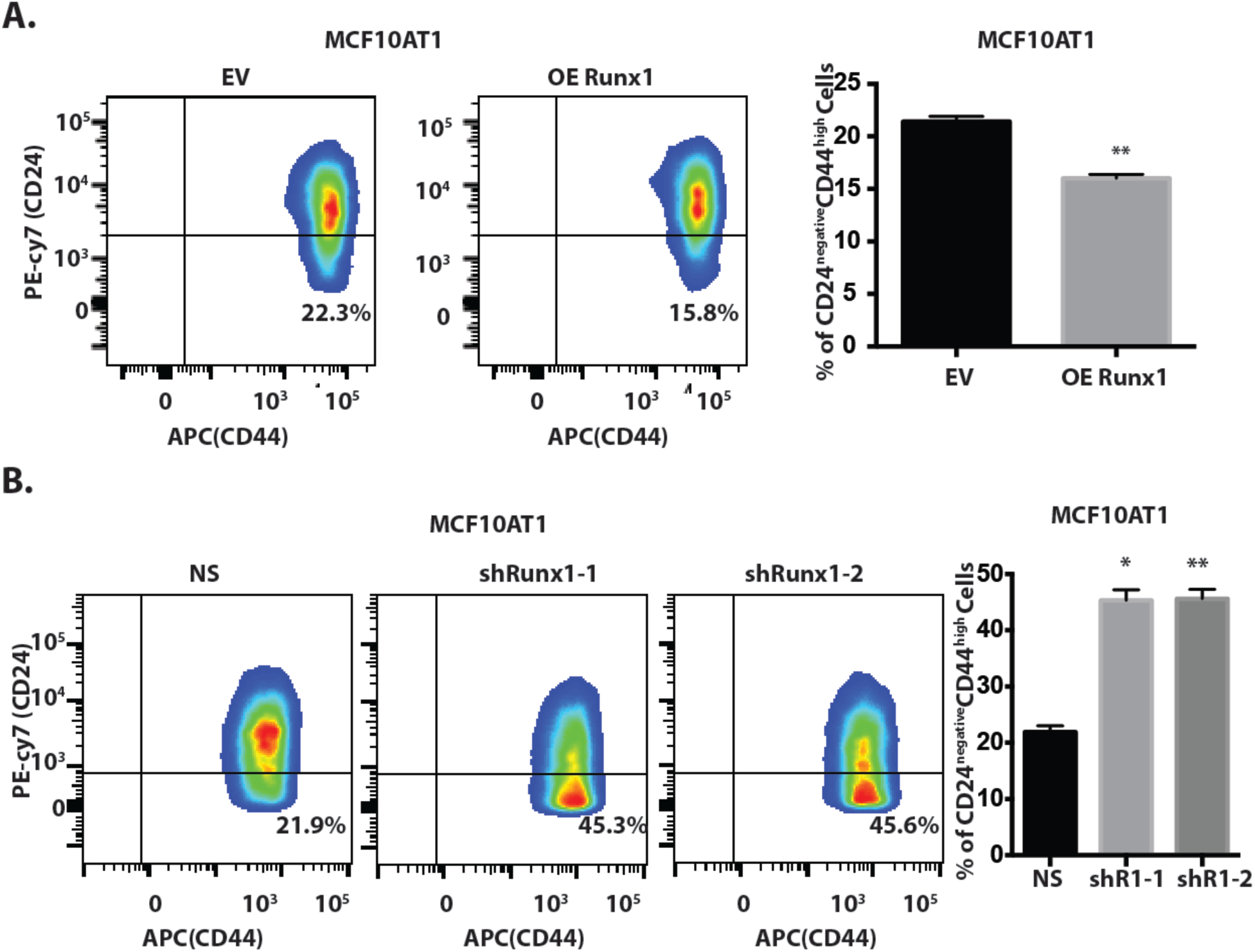
RUNX1 reduces BCSC sub-population. **(A)** Flow cytometric analysis of CD44 and CD24 expression in MCF10AT1 cells with EV or RUNX1 overexpression. Data shown represent mean ± SEM from three independent experiments. p Values were determined by student t test **(B)** Flow cytometric analysis of CD44 and CD24 expression in MCF10AT1 cells stably expressing RUNX1 or non-silencing shRNAs. Data shown represent mean ± SEM from three independent experiments. p Values were determined by student t test.

### 7. RUNX1 represses the expression of Zeb1 in breast cancer cells

In Figure 5A, we observed that decreased RUNX1 expression is coincident with activation of Zeb1 in BCSC in MCF10AT1 cells. Zeb1 is well known for its function in promoting EMT, cancer stemness and metastasis in breast cancer (43). Therefore we tested whether RUNX1 functions by negatively regulating Zeb1 expression in breast cancer cells. Zeb1 protein is down regulated when RUNX1 is ectopically expressed in MCF10AT1 cells (Fig. 7A). This RUNX1-mediated Zeb1 repression was confirmed in MCF10AT1 RUNX1 knockdown cells, where Zeb1 expression is enhanced (Fig. 7B). We did not observe RUNX1 repression of Zeb1 expression in MCF10CA1a cells, which is a consequence of very low Zeb1 mRNA levels in MCF10CA1a cells compared to MCF10AT1 cells (Supplement Figure 6). To test whether RUNX1 can directly regulate Zeb1 in MCF10CA1a cells, we performed ChIP-qPCR for RUNX1 in the Zeb1 promoter region in both MCF10AT1 and MCF10CA1a cells (Supplement Figure 7). As shown in Fig. 7C, RUNX1 directly binds to the Zeb1 promoter in the two breast cancer cell lines relative to two negative control genes ZNF333 and ZNF180. Upon RUNX1 overexpression, the binding of RUNX1 is enhanced on Zeb1 promoter, suggesting that RUNX1 has potential to direct regulate Zeb1 expression in both pre-malignant and metastatic breast cancer cell lines.

**Figure 7.**
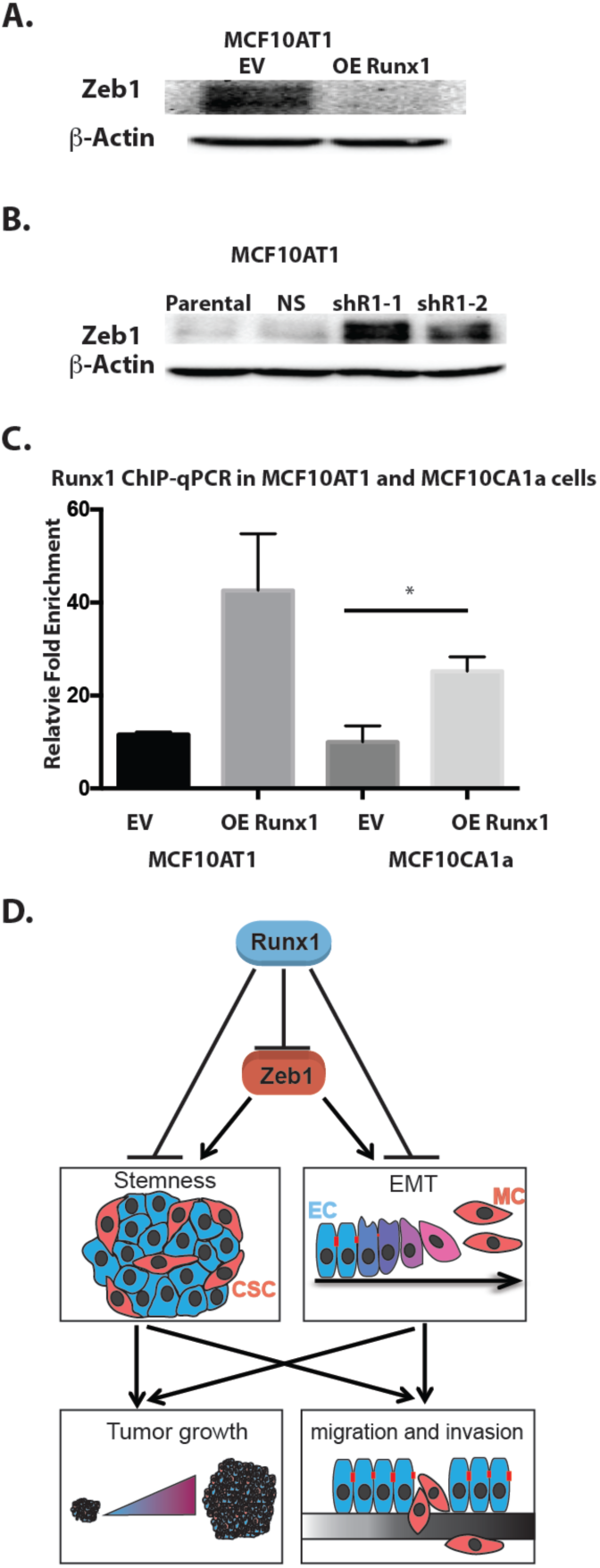
RUNX1 negatively regulates Zeb1 expression. **(A)** Western blot analyses show Zeb1 is decreased upon RUNX1 overexpression in MCF10AT1 cells. **(B)** Western blot analyses show Zeb1 is activated upon RUNX1 knockdown in MCF10AT1 cells. **(C)** ChIP-qPCR confirmation of RUNX1 occupancy at Zeb1. RUNX1 binding is increased in RUNX1 overexpression samples. Data obtained with antibodies against RUNX1 are normalized to input control and ZNF188 (NC1) and ZNF333 (NC2), which were used as the negative control as RUNX1 are predicted not to bind these genes. Data shown represent mean ± SEM from three independent experiments. p Values were determined by student t test. **(D)** Mechanism on how RUNX1 represses tumor growth in breast cancer. (EC-epithelial like cells; MC-mesenchymal-like cells).

In summary, our findings suggest that RUNX1 has tumor suppressor activity in breast cancer both *in vivo* and *in vitro*. Both EMT and cancer stem cell properties are repressed by RUNX1 in breast cancer cells. We thus conclude RUNX1-mediated repression could be through negative regulation of Zeb1 expression in breast cancer cells (Fig. 7D). Zeb1 is well known for activating both EMT and cancer stem cells in breast cancer. (43) Therefore RUNX1 indirectly represses these two cellular processes. It has been shown that Runx1 can directly repress EMT in breast cancer (32). It is possible that Runx1 can directly repress cancer stem cell phenotype in a Zeb1 independent manner (Fig. 7D). This study provides new insight into functional mechanisms of the RUNX1 transcriptional regulator in contributing to the stemness and the plasticity of breast cancer stem cells.

## Discussion

We provide multiple lines of evidence that RUNX1 has tumor suppressor activities *in vivo* by inhibiting cancer stem cell phenotypes in breast cancer. For example, RUNX1 levels are decreased in tumors grown in murine mammary fat pads. Consistent with tumor suppressor activity, RUNX1 reduces cell migration and invasion of breast cancer cells *in vitro* and tumor growth *in vivo*. RUNX1 also reduces the breast cancer stem cell population and tumorsphere formation efficiency, thus indicating that RUNX1 represses stemness properties in breast cancer. RUNX1 overexpression and knockdown studies revealed that RUNX1 mediates the mechanisms of inhibition of breast cancer stemness and tumorigenesis through repression of Zeb1 expression. Taken together, our findings provide compelling evidence that the loss of RUNX1 induces increased cancer stem cells and that RUNX1 overexpression can suppress the CSC population, which is responsible for metastasis, treatment resistance and tumor recurrence in breast cancer.

Breast cancer is ranked as the second leading cause of cancer death in women after lung cancer (44). In 2017, approximately 63,400 cases of female breast carcinoma in situ are expected to be diagnosed (45). Despite the significant advances that have been achieved in early detection and treatment of breast cancer, understanding the mechanisms of breast cancer progression and metastasis still requires intensive study. Recently, using sophisticated next-generation sequencing technology, a 40 mutation-driver gene list was generated in human breast cancer (14). RUNX1, which is often mutated in breast tumors, is one of those genes. Utilizing the TCGA clinical data sets, we found that reduced RUNX1 levels in tumor correlate with poor survival of breast cancer patients. Together these clinical findings suggest that RUNX1 may be a promising therapeutic biomarker for breast cancer screening and personalized medicine.

An unresolved question is whether RUNX1 functions as a tumor suppressor or an oncogene in breast cancer. Increasing evidence indicates that loss of RUNX1 function accompanies progression of breast cancer (29, 30, 32), supporting the concept that RUNX1 is a tumor suppressor. Clinically, RUNX1 expression is decreased in high histological grade tumors compared with low/mid grade tumors (46). In the past few years, RUNX1 loss-of-function somatic mutations have been identified in several subtypes of breast cancer (40) (47) (48). Mechanistically, loss of RUNX1 in ER+ breast cancer activates the WNT signaling pathway and ELF5 expression (29) (30) suggesting that RUNX1 represses breast cancer progression. Our previous study showed loss of RUNX1 promotes EMT in both normal and breast cancer cells indicating that RUNX1 has the potential to inhibit tumor growth (32). In this study, we clearly demonstrate using a mouse xenograft model, that the level of RUNX1 is decreased during tumor growth, and that ectopic RUNX1 expression suppresses tumor growth in the mouse mammary fat pad. Together these combined studies and our experiments establish that RUNX1 has tumor suppressor activity in breast cancer. However, we cannot rule out the possibility that RUNX1 may have other functions in breast cancer, especially in late stage disease. For example, in the MMTV-PyMT mouse model, the level of RUNX1 is positively correlated with tumor progression (35) and regulates genes promoting tumor growth in late stage MDA-MB-231 breast cancer cells (49). However in our study, we found that metastatic MCF10CA1a cells with RUNX1 overexpression formed smaller tumors in mouse mammary fat pad indicating that RUNX1 functions to reduce tumor growth. These contradictory results suggest that RUNX1 has dual functions (oncogene vs tumor suppressor) in late stage breast cancer depending on cellular context.

The tumor suppressor activity of RUNX1 in breast cancer is likely through its properties in maintaining the normal mammary epithelial phenotype. For example, loss of RUNX1 causes the cells to lose their epithelial morphology and activates mesenchymal genes in normal-like MCF10A cells (32). Furthermore, depletion of RUNX1 in ER positive luminal MCF7 breast cancer cells transforms the cells into a partial EMT state (32). It has been suggested that partial activation of the EMT promotes plasticity that allows reprogramming of the epithelial cell to acquire both migratory and stem-like features (50).

We investigated whether RUNX1 might function by suppressing Zeb1, due to its well-known activity in increasing breast cancer stemness and as a marker of EMT. Our results show that RUNX1 directly binds to the Zeb1 promoter in both MCF10AT1 and MCF10CA1a cells and that binding is enhanced upon RUNX1 overexpression. In MCF10AT1 cells, RUNX1 negatively regulates Zeb1 expression at the protein level. Together these findings indicate that the binding of RUNX1 on the Zeb1 promoter and the suppression of Zeb1 by RUNX1 reduce breast cancer stemness in cells that retain plasticity. Consistent with this conclusion, overexpressing RUNX1 in MCF10CA1a cells does not change the expression of EMT markers to the same extent that it does in premalignant MCF10AT1 cells (Fig 3A). These data and the fact that RUNX1 represses EMT in normal-like MCF10A cells (32), highlight its critical function in repressing tumor initiation and growth in early stage breast cancer. Also of significance is that overexpression of RUNX1 in MCF10CA1a cells decreased tumor growth *in vivo* and tumorsphere formation efficiency *in vitro,* suggesting that RUNX1 can have tumor suppressor activity in late stage breast cancer cells.

In summary, our findings constitute strong experimental evidence that RUNX1 functions as a tumor suppressor in breast cancer. This study provides a novel dimension to understanding how the transcriptional regulator RUNX1 contributes to the stemness and the plasticity of breast cancer stem cells. Together, these data support a central role for RUNX1 in preventing breast cancer progression. Both tight control of RUNX1 expression and RUNX1 functional integrity are required to prevent breast cancer malignancy. Consequently, clinical strategies should consider RUNX1 as a biomarker and potentially as a therapeutic candidate.

## Acknowledgments

We thank the members of our laboratory, as well as Roxana del Rio-Guerra, PhD of the UVM Flow Cytometry and Cell Sorting Facility for her help with cell sorting. The flow cytometry data we presented were obtained at the Harry Hood Bassett Flow Cytometry and Cell Sorting Facility, University of Vermont College of Medicine. This work was supported by NIH grants <u>NCI P01</u> <u>CA082834</u> (G.S.S., J.L.S.), <u>R01 CA139322</u> (G.S.S.), <u>R37 DE012528</u> (J.B.L.).

## Author contributions

D.H., J.B.L., J.L.S. and G.S.S. conceived and designed the experiments, and analyzed data. D.H., A.J.F., M.P.F. and A.L.V. performed the experiments. D.H. and K.H.F. performed animal experiments. J.R. constructed and packaged RUNX1 overexpression lentivirus. D.H., J.L.S., J.B.L. and G.S.S. wrote the manuscript.

